# Mediterranean Diet Protects Against a Neuroinflammatory Cortical Transcriptome: Associations with Brain Volumetrics, Peripheral Inflammation, Social Isolation and Anxiety

**DOI:** 10.1101/2023.11.01.565068

**Authors:** Jacob D. Negrey, Brett M. Frye, Corbin S.C. Johnson, Jeongchul Kim, Richard A. Barcus, Samuel N. Lockhart, Christopher T. Whitlow, Courtney Sutphen, Kenneth L. Chiou, Noah Snyder-Mackler, Thomas J. Montine, Suzanne Craft, Carol A. Shively, Thomas C. Register

## Abstract

**INTRODUCTION:** Mediterranean diets may be neuroprotective and prevent cognitive decline relative to Western diets, however the underlying biology is poorly understood.

**METHODS:** We assessed the effects of Western vs. Mediterranean-like diets on RNAseq generated transcriptional profiles in temporal cortex and their relationships with changes in MRI neuroimaging phenotypes, circulating monocyte gene expression, and observations of social isolation and anxiety in 38 socially-housed, middle-aged female cynomolgus macaques.

**RESULTS:** Diet resulted in differential expression of seven transcripts (FDR<0.05). Cyclin dependent kinase 14 (*CDK14*), a proinflammatory regulator, was lower in the Mediterranean group. The remaining six transcripts [i.e., “lunatic fringe” (*LFNG*), mannose receptor C type 2 (*MRC2*), solute carrier family 3 member 2 (*SLCA32*), butyrophilin subfamily 2 member A1 (*BTN2A1*), katanin regulatory subunit B1 (*KATNB1*), and transmembrane protein 268 (*TMEM268*)] were higher in cortex of the Mediterranean group and generally associated with anti-inflammatory/neuroprotective pathways. *KATNB1* encodes a subcomponent of katanin, important in maintaining microtubule homeostasis. *BTN2A1* is involved in immunomodulation of γδ T-cells which have anti-neuroinflammatory and neuroprotective effects. *CDK14*, *LFNG*, *MRC2,* and *SLCA32* are associated with inflammatory pathways. The latter four differentially expressed cortex transcripts were associated with monocyte transcript levels, changes in AD-relevant brain volumes determined by MRI over the course of the study, and social isolation and anxiety. *CDK14* was positively correlated with monocyte inflammatory transcripts, changes in total brain, gray matter, cortical gray matter volumes, and time alone and anxious behavior, and negatively correlated with changes in total white matter and cerebrospinal fluid (CSF) volumes. In contrast, *LFNG*, *MRC2*, and *SLCA32* were negatively correlated with monocyte inflammatory transcripts and changes in total gray matter volume, and positively correlated with CSF volume changes, and SLCA32 was negatively correlated with time alone.

**DISCUSSION:** Collectively, our results suggest that relative to Western diets, Mediterranean diets confer protection against peripheral and central inflammation which is reflected in preserved brain structure and behavior.

## Introduction

An increasing literature suggests that diet composition may effect mental health, neurobiological aging, the rate of cognitive decline with aging, and the risk of neurodegenerative disorders including Alzheimer’s disease (AD) [1–3]. The Mediterranean-style diet (hereafter, “Mediterranean diet”), which is largely plant-based and rich in healthy oils, complex carbohydrates, and dietary fiber, is associated with increased longevity [4] and quality of life [5], lower rates of psychiatric [3, 6] and neurodegenerative disease, and slower cognitive aging [7, 8]. as well as reduced risk of other chronic health conditions [9–12]. A short-term randomized clinical trial has shown positive effects of a Mediterranean-like diet on CSF markers relevant to AD pathology [13]. Conversely, consumption of a so-called “Western” diet, which is high in saturated fats and simple sugars [14, 15], has been shown to increase systemic inflammation [16] and have detrimental neurobiological effects. Short term consumption of very high fat diets (60-75% fat) impairs attention and memory in clinical and preclinical studies [17]. Observational studies show associations of Western diet consumption with increased risk of depression, anxiety [6, 18] and faster cognitive decline with aging [19, 20]. Furthermore, Western diet consumption is associated with brain-related changes, such as blood-brain barrier disruption and neuroinflammation, which could contribute to neurocognitive decline and AD [21, 22]. However, conflicting findings have been reported. Of 10 of the most comprehensive trials of cognitive effects of the Mediterranean diet reviewed by Morris et al [23], only four reported clear protective effects against cognitive decline or dementia [24–27]. Likewise, a recent evaluation of the association between adherence to the MIND (Mediterranean-Dietary Approaches to Stop Hypertension Diet Intervention for Neurodegenerative Delay) diet and cognitive health in the UK Biobank reported no association with better cognitive test scores and lower dementia risk only in women [28]. These conflicting results may be due to how Mediterranean diet consumption was measured [23]. The majority of these data derive from observational studies that rely on self-reported diet consumption and may be confounded by other factors which correlate with diet consumption (e.g., activity levels [29]) and are known to impact neurocognitive outcomes. Consequently, experimental studies testing neurobiological effects of diet composition are essential to determine whether diet patterns directly impact the brain.

Given genetic, physiological, and social similarity to humans, nonhuman primates are useful models for aging and age-related disease and provide important opportunities for studying the molecular mechanisms of disease. Macaques (*Macaca* spp.) exhibit neurobiological aging and cognitive decline, and develop early AD-like neuropathologies including amyloid accumulation and tauopathies [30, 31].

In a randomized preclinical trial, our group has shown that Mediterranean diet consumption by female macaques protects against overeating, obesity, hepatosteatosis, insulin resistance [32], and increased hypothalamic-pituitary-adrenal and sympathetic nervous system activity [33] compared to Western diet consumption. In this same study, we also assessed the effects of diet on longitudinal structural magnetic resonance imaging (MRI) measures of neuroanatomy and found that global brain volumes changed in response to Western but not Mediterranean diets [34]. Monkeys fed the Western diet showed increased total brain (TBV) and gray matter (GM) volumes and decreased volumes of cerebrospinal fluid (CSF) and white matter (WM) after 31 months of experimental diet consumption. The Western diet group also had increased cortical thickness in multiple regions in the temporal and parietal lobes relevant to AD [35]. Given these observations, we hypothesized that diet-related increases in total brain volume and GM thickness as well as decreases in WM volume were likely due to Western diet-induced neuroinflammation [34]. This hypothesis was spurred, in part, by clinical data from the Alzheimer’s Disease Neuroimaging Initiative (ADNI) trial indicating biphasic changes in brain volumes over time, with increased gray matter volumes and cortical thickening early in the AD disease process (asymptomatic amyloidosis) followed by losses and AD pathology and neurodegeneration increases [36]. The investigators hypothesized that these increased volumes may reflect neuroinflammation [36–38], which, with time, results in cortical thinning, a predictor of increased AD risk [39]. Consistent with this hypothesis, in our NHP study, circulating monocytes in the Western group were polarized towards a pro-inflammatory phenotype, relative to those from the Mediterranean group, characterized by increased transcripts for IL6, IL1α, NFKB1, and NFKB2, key transcriptional regulators of inflammatory pathways [40]. Likewise a beneficial circulating inflammatory profile is associated with Mediterranean diet consumption in human studies [41]. Furthermore, in our NHP trial the experimental diets induced rapid and persistent changes in a suite of behaviors. Monkeys fed the Western diet spent more time alone and displayed more anxiety behavior, whereas those fed the Mediterranean diet spent more time resting, attentive, and in body contact with groupmates. Finally, the indices of peripheral inflammation in this study were associated with social isolation and anxiety, behaviors thought to be associated with neuroinflammation [42].

In this, the first study to investigate differential effects of Western vs. Mediterranean diets on the brain transcriptome, we sought to identify molecular mechanisms underlying diet-induced neuroanatomical changes by testing if diet composition altered gene expression in the temporal lobe cortex of socially housed, female long-tailed macaques (*Macaca fascicularis*). We focused on females because there is a paucity of data from female animal models, and because many brain diseases are more prevalent in females than males. For example, two-thirds of the Alzheimer’s disease burden is in women [43]. We focused on middle-age because that is the earliest period in which the neuropathological trajectory that culminates in cognitive decline and neurodegeneration has been observed; thus identification of targets for prevention is needed at this life stage. We focused on the temporal cortex because this region is among the first to exhibit neuropathological signs of AD [44, 45]. We first identified differentially expressed genes (DEGs) between the two diets and then examined relationships between gene expression and changes in brain volumes and cortical thicknesses determined by MRI between baseline and after 31 months of experimental diet. Based on our prior findings [33, 34, 40, 46, 47], we hypothesized that monkeys fed a Western diet would exhibit transcriptomic signals of inflammation that would be associated with peripheral inflammation, changes in brain volumes and behavior, potentially illuminating molecular targets for intervention. Notably, we explore transcriptomic values in the context of MRI-derived brain values, which provide clear translational value. We also examined associations of temporal cortex transcript levels with peripheral monocyte transcripts, and with levels of anxiety and social isolation.

## Methods

### Ethics Statement

All experimental procedures complied with National Institutes of Health Guide for the Care and Use of Laboratory Animals (NIH Publications No. 8023, revised 1978), state and federal laws, and were approved by the Animal Care and Use Committee of Wake Forest School of Medicine.

### Subjects

Details of the experiment and subjects have been previously described [33, 34, 46, 48]. Briefly, adult female cynomolgus (a.k.a. long-tailed) macaques (*M. fascicularis*) aged 11 to 13 years (as estimated from dentition) were obtained from SNBL USA (Alice, Texas) and quarantined for one month in single-cages, during which time subjects had both visual and auditory contact with each other. Following quarantine, monkeys were transferred to social groups of n=4-5 each and housed in indoor enclosures (3m X 3m X 3m) with natural light exposure, 12h/12h light-dark cycles, and water available *ad libitum*. All individuals were fed monkey chow (LabDiet) during a 7-month baseline period prior to the experimental dietary intervention.

### Experimental Procedure

After the 7-month baseline phase, the monkeys were assigned to one of two diet groups: Western or Mediterranean. The experimental groups were balanced regarding several relevant subject characteristics, including body composition (body weight and body mass index), circulating basal cortisol, and fasting triglyceride concentrations using stratified randomization [32]. Experimental diets were developed and prepared at the Wake Forest School of Medicine Comparative Medicine Diet Lab and fed by animal care staff unassociated with the research project. Samples of all diet batches were kept. Batch analyses demonstrated consistency over time. Diets were balanced for macronutrients and cholesterol content, but differed in sources of protein, fat, and carbohydrates. Diet compositions are detailed in **Supplementary Table 1** and the complete ingredient list may be found in [32]. The Western diet, which was designed to resemble that consumed by middle-aged women (40-49 years) living in the USA [14], featured primarily animal-derived fats and proteins, and a relatively high content of simple sugars and salt [15]. In contrast, the Mediterranean diet featured plant-based fats and proteins, resulting in a diet high in monounsaturated fats, complex carbohydrates, and fiber, and low in sodium and refined sugars [15, 49]. Importantly for inflammation, the Western diet had relatively higher amounts of omega-6 fatty acids, whereas the Mediterranean diet was higher in omega-3 content. English walnut powder and extra virgin olive oil, as given to participants in the PREDIMED study [50], were among the chief dietary components of the Mediterranean diet. Thirty-eight monkeys (21 Western, 17 Mediterranean) completed the experiment and were subsequently assessed in this study.

### Social Isolation and Anxiety Behavior Observations

The behavioral characterization of these animals has been described in-depth [51]. Briefly, behavioral data were collected weekly during two 10-min focal observations [52], balanced for the time of day for 24 consecutive months beginning the third month of the treatment phase (approximately 200 behavior samples/monkey, mean = 31.0 h/monkey, and 1178 observation hours total). Inter-rater reliability was maintained at ≥93%, and no other research activities were ongoing during behavior observations. Since we previously determined that the Western diet increased social isolation and anxiety, we focused on these behaviors here. Social isolation was operationally defined as the rate of times a monkey spent alone out of arm’s reach of another monkey or in anxiety related behavior. Anxiety was operationally defined as the frequency self-grooming and scratching [53–57].

### Magnetic Resonance Imaging

Neuroimaging methods used in this study have been previously described [34]. Briefly, T1-weighted structural MRI scanning was performed at baseline and again at 31 months at the MRI facility at Wake Forest University School of Medicine using a 3T Siemens Skyra scanner (Siemens, Erlangen, Germany) with a 32-channel head coil (Litzcage, Doty Scientific, SC). Brain tissue from scanned images was segmented into three main types: gray matter, white matter, and cerebrospinal fluid. We then used the parcellation map to determine volumes for the following regions of interest (ROIs): total gray matter (tGM), cortical gray matter (cGM), white matter (WM), cerebrospinal fluid (CSF), and total brain volume (TBV (tGM + WM)). The cortical thicknesses were determined using T1w MR image and the cortical GM tissue segmentation in the subject’s native space. We also generated volume- and thickness-based AD temporoparietal meta-ROIs using the AD clinical literature as a template [37] using a previously described method [34]. Each meta-ROI represented the following ROIs (as illustrated in **Figure 1**): the angular gyrus, the inferior, middle, and superior temporal gyrus, entorhinal cortex, fusiform cortex, supramarginal gyrus, precuneus, and parahippocampus. A detailed description of imaging methods is available in the **Supplementary Information**.

**Figure 1.**
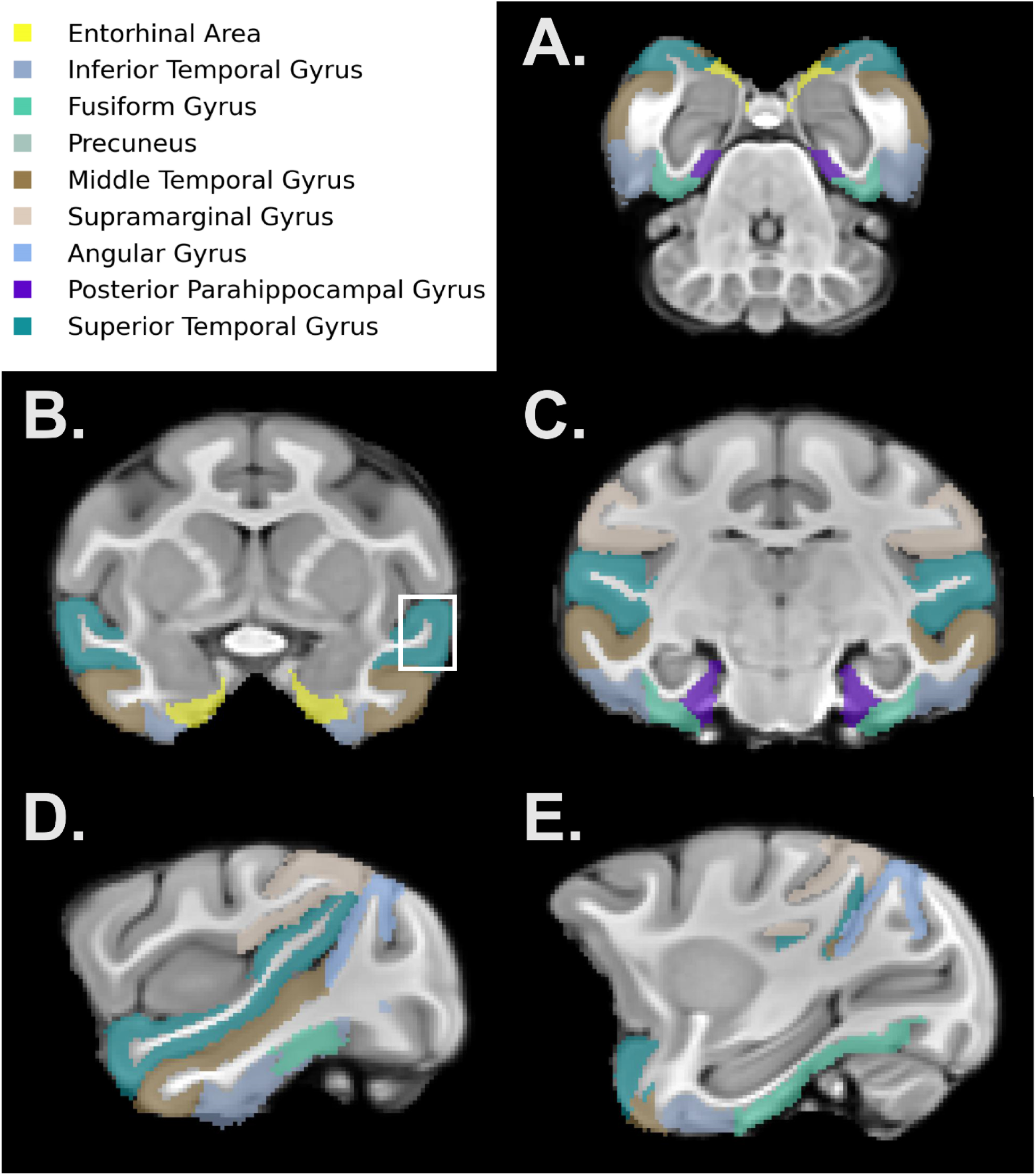
Parcellation map of macaque brains illustrating regions included in Alzheimer’s disease meta-regions of interest. Images are provided in axial (panel A), coronal (panels B and C), and sagittal (panels D and E) cross-sections. The white box (panel B) indicates a box approximating the region from which gray matter was collected for transcriptomic profiling.

### Peripheral Monocyte Transcriptomics

Details regarding isolation and purification of CD14+ monocytes, RNAseq analyses, and subsequent findings have been previously published [40]. Briefly, CD14+ monocytes were purified from PBMCs using positive selection with Miltenyi anti-CD14 magnetic beads. RNA was extracted from monocytes using the AllPrep DNA/RNA Mini Kit (Qiagen, Inc, Hilden, Germany), and quantified using a NanoDrop spectrophotometer and Agilent 2100 Bioanalyzer with RNA 6000 Nano chips (Agilent Technology, Inc, Santa Clara, CA). RNA libraries were prepared for sequencing by the Cancer Genomics Shared Resource (Wake Forest University School of Medicine, Winston-Salem, NC) using the TruSeq-stranded total RNA kit (Illumina), which includes a ribosomal depletion step. RNA-seq libraries were sequenced using single-end 76 bp reads on an Illumina NextSeq 500 to an average read depth of 34.5 million reads per sample (range 25.9–41.6 million reads). Reads were mapped to the M. fascicularis reference genome (Macaca_fascicularis_5.0, v 93, Ensembl using HiSat2 and then converted to a sample-by-gene read count matrix using featureCounts (Liao et al., 2014) (median = 38.0%; range 24.5–50.4% of reads mapped to exons). Sample processing order was randomized and where possible all samples were manipulated simultaneously so as to avoid introducing batch effects. Genes with low expression (median reads per kilobase per million reads mapped < 1) were removed, which resulted in 12,240 genes for downstream analyses. Read counts were normalized using the voom function of the R package limma (Ritchie et al., 2015). The final sample size was 35 monkeys (n = 20 fed the Western diet, n = 15 Mediterranean diet). To control for batch effects related to RNA quality and monocyte purity, we calculated the residual gene expression from a model of normalized gene expression as a function of CD14 expression, CD3 expression, RNA integrity, and RNA concentration. These residual gene expression values were used for all subsequent analyses.

### Tissue Collection and Processing

Following the 31-month exposure to the experimental diets, monkeys were anesthetized with pentobarbital (30-50 mg/kg) to obtain a surgical plane of anesthesia, followed by exsanguination and perfusion with ice cold saline, according to the guidelines of the American Veterinary Medical Association’s Panel on Euthanasia. To provide high quality samples for transcriptomic analyses, we implemented short post-mortem intervals (i.e., from euthanasia to brain preservation). The brain was hemisected and blocked into coronal slabs 4mm in thickness; tissue collected for transcriptomic analyses was covered with finely crushed dry ice and frozen for at least 20 minutes, then placed in a vacuum-sealed sample bag and stored in a - 80°C freezer. Detailed examination of brain showed generally normal brain architecture and absence of amyloid plaques or tauopathies, consistent with the age of the monkeys. The tissue sampled for this study is indicated in **Figure 1**.

### RNA-seq and Bioinformatics

RNA isolation and sequencing were performed by the Wake Forest School of Medicine Cancer Genomics Shared Resource which were blinded to treatment. RNA was isolated from temporal cerebral cortex samples using the Zymo Quick-DNA/RNA Miniprep Plus Kit (Zymo, Irvine, CA, USA) and purified using the RNA Clean and Concentrator-5 kit (Zymo). All RNA integrity number (RIN) scores were > 7.7, indicating high integrity. Bulk RNA-seq was then performed using an Illumina NovaSeq 6000 (single end, 100bp). We subsequently aligned sequenced reads to the long-tailed macaque genome (Macaca_fascicularis_6.0) using the STAR alignment software [58].

Differential gene expression (DGE) analysis was performed in R version 4.1.1 [59] using the *pqlseq* function in package “PQLseq” [60], which controls for genetic relatedness between subjects while performing DGE analysis within the framework of a generalized linear mixed model. We first filtered genes exhibiting low reads per kilobase of transcript per million mapped reads (RPKM), leaving 10,684 genes for DGE analysis. To determine relatedness, we used *bcftools mpileup* [61] to infer SNP genotypes from RNA-seq reads, as previously described [40]. Based on this approach we found 1 full sib pair and 9 other pair combinations which were half-sibs. We modeled the expression of each gene as a function of diet (Western or Mediterranean) and RIN (to control for any effects of RNA quality on expression). Given that social status [62, 63] and estradiol [64] exert neurological effects in NHPs, we also included subject social status (relative dominance rank) and serum estradiol levels at time of necropsy as covariates. Genes were considered differentially expressed if they passed a Benjamini-Hochberg false discovery rate threshold of 0.05.

### Identifying Biological Pathways and Causal Networks

Biological processes and regulatory networks exhibiting diet-specific activation patterns were investigated using Ingenuity Pathway Analysis (IPA; Qiagen, Hilden, Germany). Given the limited number of DEGs identified at a false discovery rate (FDR) cutoff of 0.05, for exploratory analyses we broadened our criteria to include genes with an FDR<0.2 (n=312), following prior work [40]. First, we extracted canonical pathways associated with the gene set, retaining all pathways for which p<0.05. We then analyzed causal networks to identify “master” regulators associated with the gene set; we retained only endogenous regulators, discarding all exogenous substances (e.g., drugs, chemical reagents). To determine whether causal networks and associated regulators were significantly activated or inhibited, we employed activation Z-scores, which indicate the confidence that a regulatory network is activated or inhibited based on transcript levels within a gene set. We used thresholds of >2 (activated) and <-2 (inhibited) and discarded regulators with non-significant Z-scores (i.e., Z-score between 2 and −2).

### Correlations between Temporal Cortex Gene Expression, Brain Volumes, Peripheral Monocyte Transcripts, and Behaviors

We assessed relationships between transcript levels from genes that were differentially expressed by diet and circulating monocyte gene expression, changes in neuroanatomy, and behavior using the *corr.test* function in R. For more information, see the **Supplementary Information**.

## Results

### Temporal Cortex Gene Expression Varied by Diet

Seven genes were significantly differentially expressed between the two diets (FDR<0.05; Figure 2). Cyclin dependent kinase 14 (*CDK14*), was downregulated in the Mediterranean cohort relative to the Western cohort (β=-0.087, SE=0.021, p=2.85E-05, FDR= 0.044). The remaining six genes were upregulated in the Mediterranean cohort relative to the Western cohort: butyrophilin subfamily 2 member A1 (*BTN2A1*; β=0.172, SE=0.041, p=2.45E-05, FDR= 0.044), katanin regulatory subunit B1 (*KATNB1*; β=0.118, SE=0.028, p=2.68E-05, FDR=0.044), beta-1,3-N-acetylglucosaminyltransferase lunatic fringe, or “Lunatic Fringe” (*LFNG*; β=0.253, SE=0.053, p=1.93E-06, FDR=0.021), mannose receptor C type 2 (*MRC2*; β=0.324, SE=0.073, p=9.58E-06, FDR=0.032), solute carrier family 3 member 2 (*SLC3A2*; β=0.119, SE=0.027, p=1.21E-05, FDR=0.032), and transmembrane protein 268 (*TMEM268*; β=0.157, SE=0.035, p=7.51E-06, FDR=0.032). Of these seven DEGs, Ingenuity Pathway Analysis identified four as having known causal relationships or associations with cytokines: *CDK14*, *LFNG*, *MRC2*, and *SLC3A2* (**Figure 3**).

**Figure 2.**
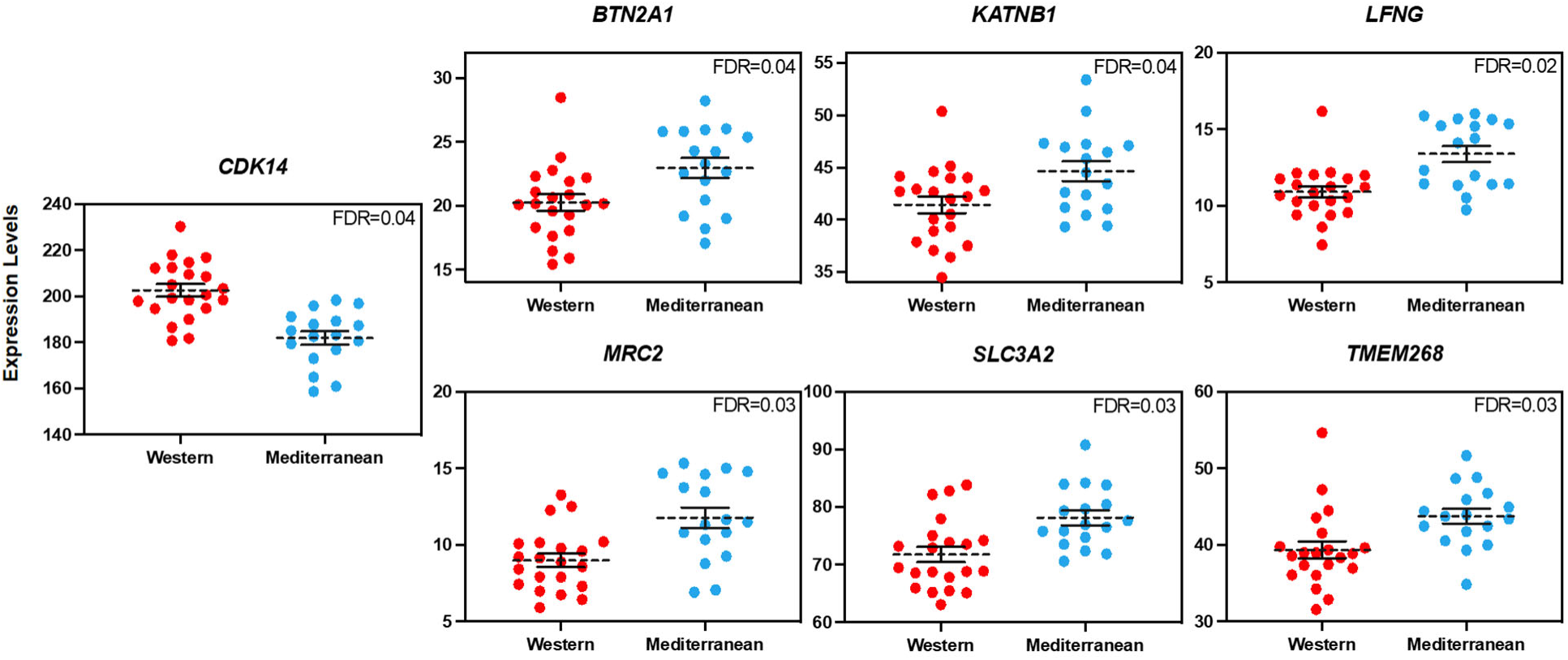
Transcript levels (TMM normalized) for seven genes differentially expressed by diet: Cyclin dependent kinase 14 (CDK14), butyrophilin subfamily 2 member A1 (BTN2A1), katanin regulatory subunit B1 (KATNB1), “Lunatic Fringe” (LFNG), mannose receptor C type 2 (MRC2), solute carrier family 3 member 2 (SLC3A2), and transmembrane protein 268 (TMEM268).

**Figure 3.**
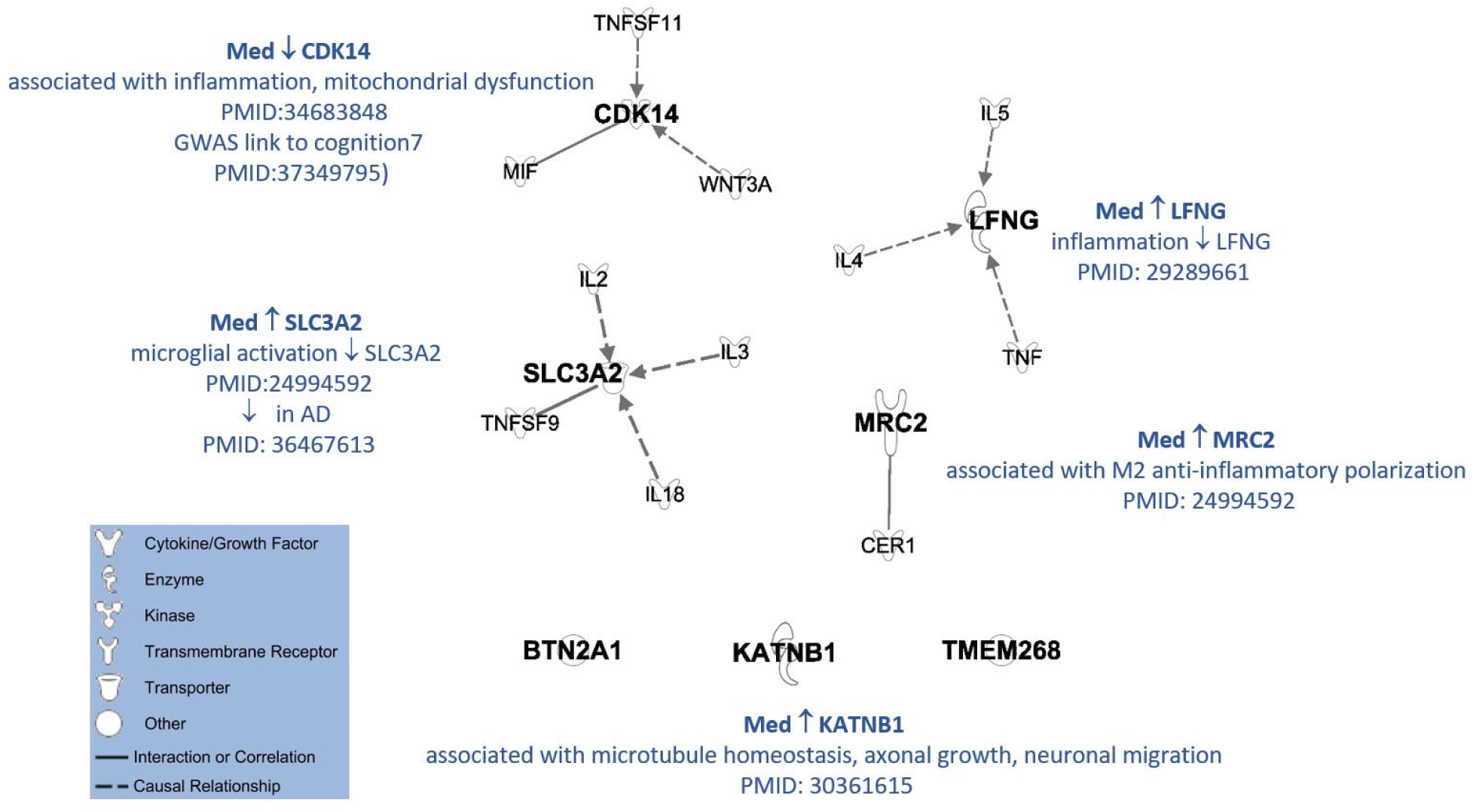
Associations between differentially expressed genes (bold, italicized font) and cytokines determined using Ingenuity Pathway Analysis. Solid and dashed lines indicate direct and indirect interactions, respectively. Arrows indicate directional relationships.

To determine whether diet effects were driven by differences in cell type proportions, we estimated cell type proportions for each sample (Supplementary Information). Cell type biases for each sample are available in **Supplementary Table 2**. We did not find diet-based differences in the estimated proportion of any of six brain cell types (**Supplementary Table 3**). Furthermore, Wilcoxon rank sum tests of the 7 DEGs indicated that diet-based differences persisted even after correcting transcript levels for cell type proportions (**Supplementary Table 4**).

At a relaxed FDR threshold (0.2), we found 312 diet-associated DEGs (**Supplementary Table 5**). These genes were involved in: (1) DNA Double-Strand Break Repair by Non-Homologous End Joining, (2) Phosphatase and Tensin Homolog (PTEN) Signaling, (3) Ciliary Neurotrophic Factor (CNTF) Signaling, (4) Polyamine Regulation in Colon Cancer, and (5) Tumor Microenvironment Pathway. (All associated pathways are listed in **Supplementary Table 6.**) The top five master regulators identified by IPA included (1) Sp3 Transcription Factor, (2) Chymase 1 (CMA1), (3) LXR ligand-LXR-Retinoic acid-RXR, (4) Transcriptional Repressor GATA Binding 1 (TRPS1), and (5) transforming growth factor beta 1 (TGFB1).

*SP3* (Z-score=2.26), *CMA1* (Z-score=3.46), LXR ligand-LXR-Retinoic acid-RXR (Z-score=2.24), and *TGFB1* (Z-score=2.83) were more highly expressed in the Mediterranean cohort, whereas *TRPS1* was lower (Z-score=-2.95). A complete list of causal networks with associated master regulators is available in **Supplementary Table 7**. IPA identified four of the seven DEGs to have known causal relationships or associations with cytokines (**Figure 3**). To address our central hypothesis that monkeys fed a Western diet would exhibit transcriptomic signals of inflammation that would be associated with peripheral inflammation, changes in brain volumes, and behavior, we focused on these four DEGS in the following correlational analyses with brain structure, circulating monocyte gene expression, and behavior.

### Temporal Cortex Gene Expression was Associated with Circulating Monocyte Gene Expression

In order to explore potential relationships between peripheral and central gene expression, we assessed relationships of differentially expressed transcripts in temporal cortex at necropsy with circulating CD14+ monocyte transcript levels midway through the project. Patterns of gene expression observed in monocytes predicted patterns observed in the brain, with consistent anti-inflammatory effects of the Mediterranean Diet relative to the Western Diet (**Figure 5**). Monocyte transcripts associated with inflammation, including inflammatory components of the family of genes represented in the CTRA (the constitutive transcriptional response to adversity), NFKB1, NFKB2, and IL6, were all positively associated with transcript levels of the pro-inflammatory gene CDK14 in the temporal cortex. In general, these genes were all negatively correlated with anti-inflammatory transcripts LFNG, SLC3A2, and MRC2 in the temporal cortex (11 of 12 transcripts p< 0.10, 8 of 12 transcripts p<0.05).

### Temporal Cortex Gene Expression was Associated with Changes in Brain Structure

To determine whether expression levels for the four DEGs known to have causal relationships or associations with cytokines were reflected variation in brain structure, we ran a series of Pearson’s correlations in which global brain volumes and temporoparietal meta-ROIs were functions of gene expression levels (**Figure 4**). On the basis of uncorrected p values, *CDK14* transcript levels were significantly positively correlated with percent change in total brain volume, total gray matter volume, cortical gray matter volume, thickness of the temporoparietal meta-ROI in both hemispheres, and volume of the temporoparietal meta-ROI in the right hemisphere, and negatively correlated with percent change in global white matter and CSF volumes (all p<0.05, **Figure 4**). In contrast, *LFNG*, *MRC2*, and *SLC3A2* transcript levels were significantly negatively correlated with percent change in total brain volume, total gray matter volume, and thickness and volume of the temporoparietal meta-ROI in the right hemisphere, and positively correlated with percent change in global CSF volume (p<0.05, **Figure 4**). *SLC3A2* transcript levels were also positively correlated with percent change in global white matter volume, and *MRC2* and *SLC3A2* were negatively correlated with thickness of the temporoparietal meta-ROI in the left hemisphere (**Figure 4**). We also analyzed each diet cohort independently to determine whether patterns were attributable to diet composition or upheld within each diet. When doing so, we found no significant correlations within the Mediterranean cohort (**Supplementary Table 8**) but did within the Western cohort; *MRC2*, in particular, was significantly correlated with seven of the nine MRI measures in the Western subset (**Supplementary Table 9**).

**Figure 4.**
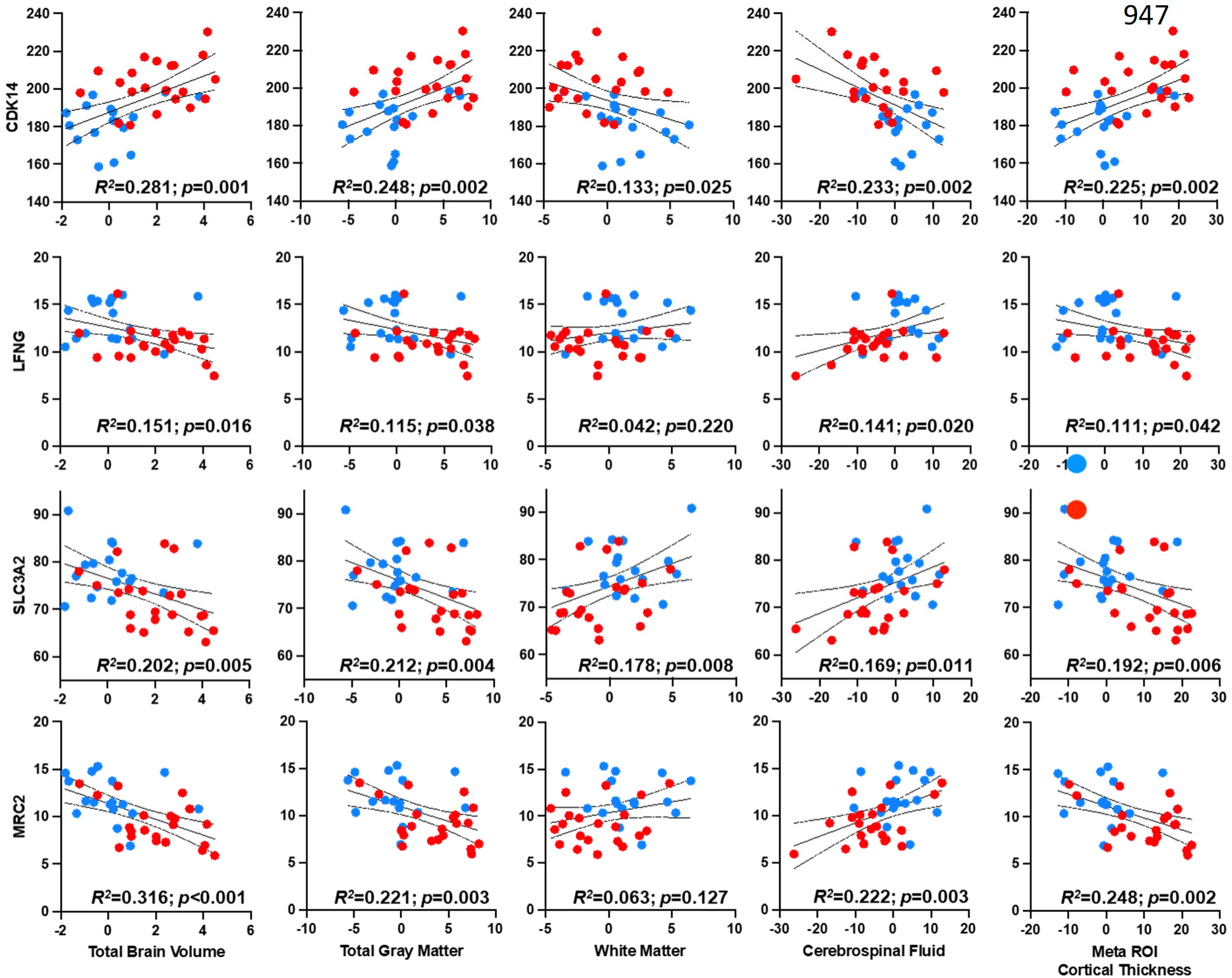
Scatterplots of transcript levels (TMM normalized; y axis) for genes with known inflammatory associations (CDK14, LFNG, SLC3A2, and MRC2) versus percent changes in brain tissue / fluid volumes (x axis) from baseline to end of experimental phase.

**Figure 5.**
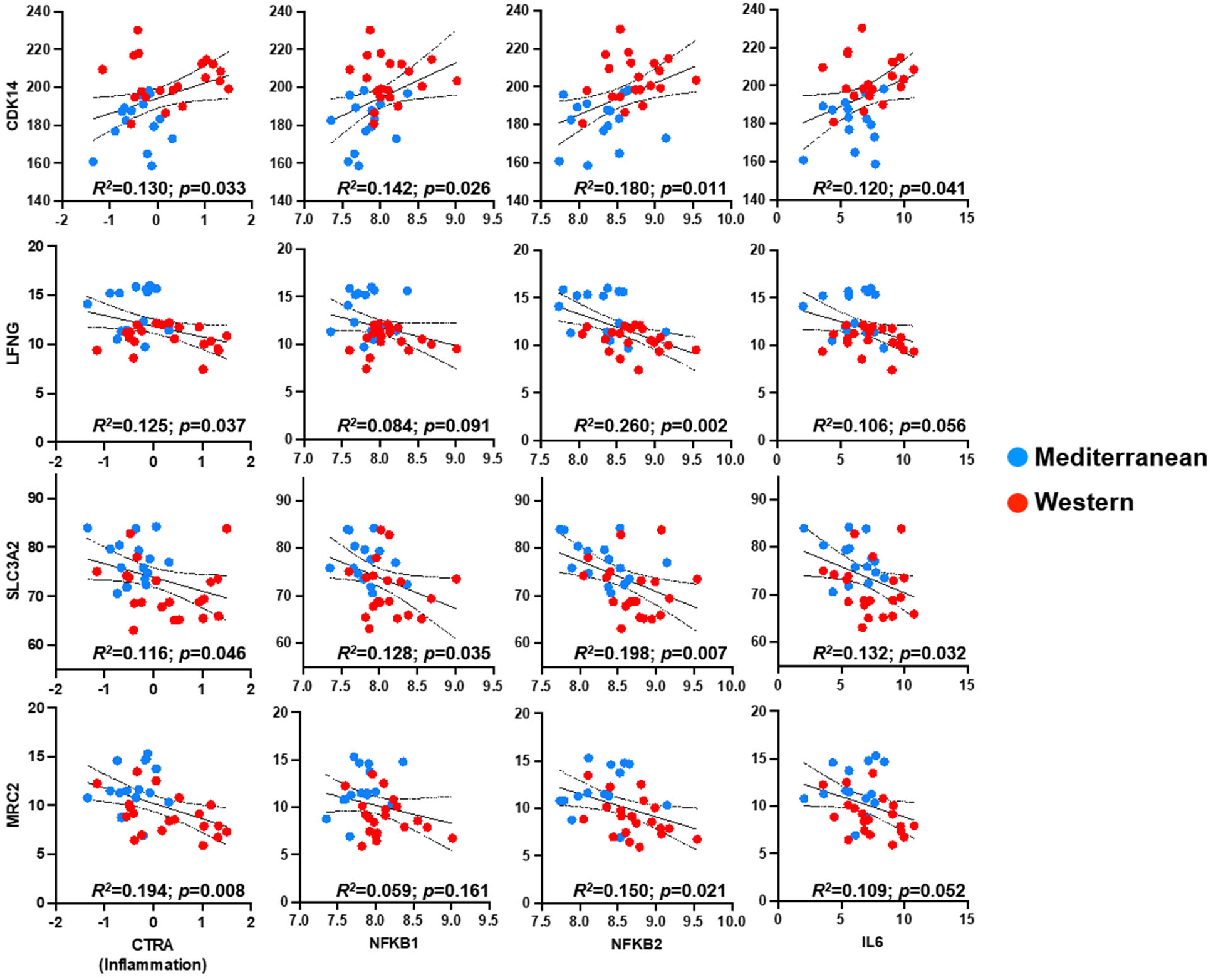
Relationships of differentially expressed transcripts in temporal cortex at necropsy with circulating CD14+ monocyte transcript levels midway through the project.

### Temporal Cortex Gene Expression Was Associated with Behavior

Temporal cortex transcript levels of *CDK14* were positively correlated with time spent alone and with anxious behavior. In contrast, levels of the anti-inflammatory transcript for *SLCA32* were negatively correlated with time spent alone (**Figure 6)**.

**Figure 6.**
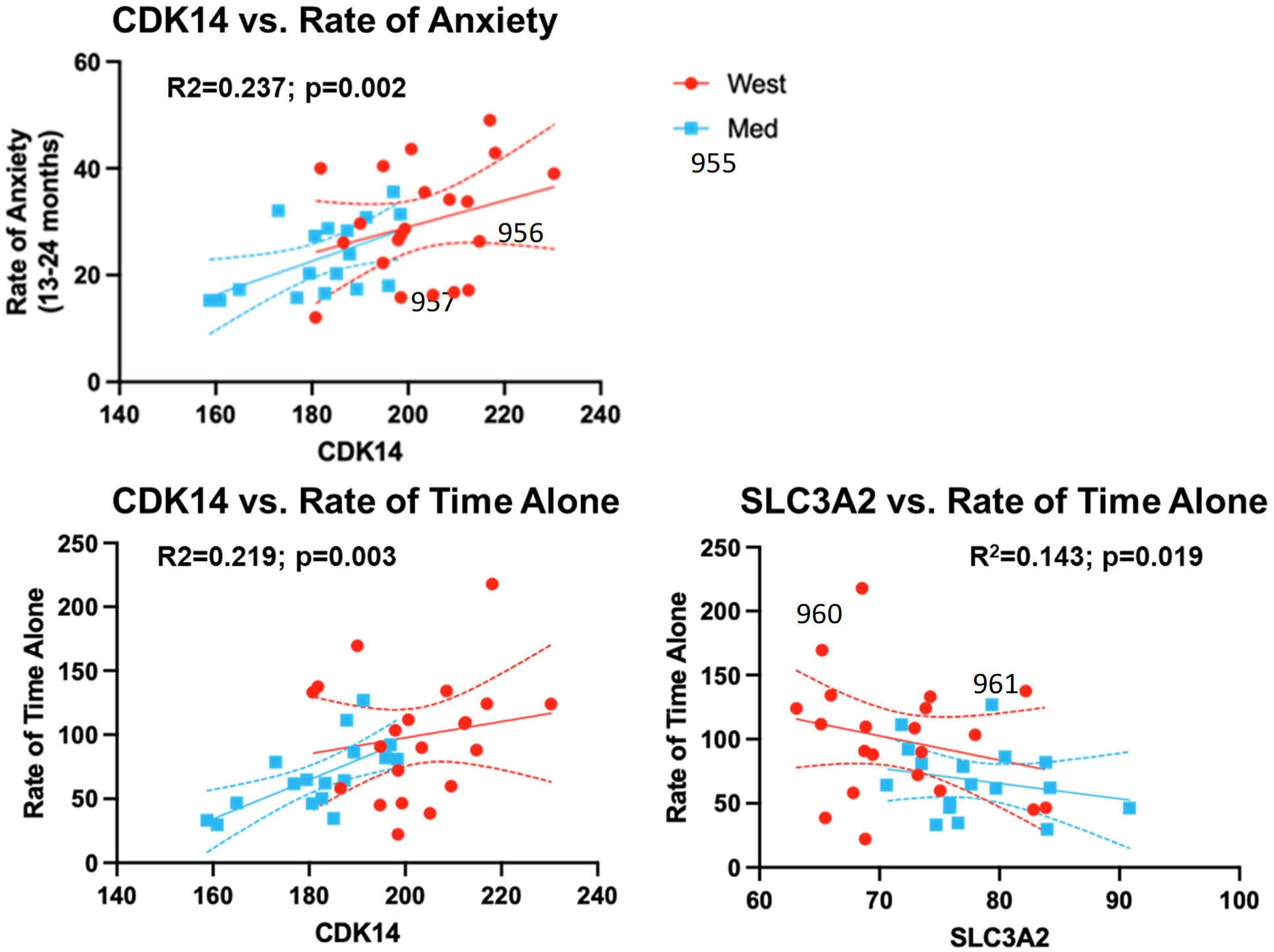
Lateral temporal neuroinflammatory transcriptome associations with anxiety and time spent alone. Temporal cortex CDK14 transcripts were positively associated with rates of anxiety and time spent alone, while cortex SLC3A2 was inversely associated rate of time spent alone.

## Discussion

We observed differences in transcript levels in the temporal cortex of macaques fed Western versus Mediterranean diets. Four of these DEGs, known to have causal relationships or associations with cytokines, showed consistent, significant correlations with circulating monocyte gene expression, changes in brain volume and temporoparietal cortical thicknesses, and social isolation and anxious behavior. These data support our previous hypothesis that diet-related changes in neuroanatomy – i.e., increases in total brain volume and cortical thickness as well as decreases in white matter volume [34, 35]—may reflect Western diet-induced inflammation [34], whereas relative stability in such neuroanatomical profiles in the Mediterranean diet consumers may reflect resilience and heathy aging during middle-age. Furthermore, these data provide links between cortical inflammatory profiles, peripheral monocyte programming, and behavioral phenotypes. Understanding patterns during this stage of life may be clinically relevant given the extended preclinical phase of neurodegenerative disorders like AD.

Our transcriptomic results support the hypothesis that the Western diet promoted and the Mediterranean diet protected against neuroinflammation, as several DEGs identified in this study are explicitly associated with inflammation. The single gene found to be increased in the Western cohort, cyclin dependent kinase 14 (*CDK14*), regulates the Wnt signaling pathway [65], an important inflammatory modulator [66] implicated in AD progression [67]. In clinical applications, *CDK14* has been targeted for its role in tumorigenesis [68]. *CDK14* contributes to cell cycle regulation [69], and its inhibition effectively limits cell proliferation [70]. In a multiomics study to identify potential targets for AD, assessment of transcriptional profiles from 5 different studies of AD patients and controls identified a set of common DEGs; CDK14, among others, was identified as being co-expressed in a brain specific manner with this set of genes [71].

Neuropathological or neuroinflammatory actions by *CDK14* are further implicated in our dataset by its correlations with brain volumes: this gene was positively correlated with total brain volume, total gray matter volume, and temporoparietal meta-ROI thicknesses, and negatively correlated with white matter and CSF volumes, suggesting Western diet-induced inflammation in gray matter. Such inflammation in middle-age may predispose individuals for developing AD pathology, resulting in reduced hippocampal and AD-signature region volumes later in life [72]. Previously we showed that the indices of peripheral inflammation in this study were associated with social isolation and anxiety, behaviors thought to be associated with neuroinflammation [42]. Here we show that brain CDK14 transcript levels were positively associated with anxiety behavior and time spent alone consistent with previous associations of Western diet consumption with increased risk of anxiety [6].

The inflammation hypothesis is further supported by other transcripts which were significantly lower in the Western cohort. “Lunatic fringe” (*LFNG*) regulates Notch signaling [73], which is in turn an indispensable regulator of neural progenitor cell renewal [74, 75]. Experimental inflammation stimulated by tumor necrosis factor alpha causes downregulation of *LFNG* in rodent brain endothelial cells [76], which may represent an important target by which inflammatory mediators disrupt the blood-brain barrier [77] and promote pathology of the central nervous system (CNS) [76, 78]. Furthermore, LFNG was negatively associated with total brain volume, total gray matter volume, and temporoparietal meta-ROI thicknesses, and positively associated with CSF volume.

Similarly, mannose receptor C type 2 (*MRC2*), which encodes a receptor expressed on the surface of fibroblasts, M2 macrophages and microglia [79, 80], was also upregulated in the Mediterranean cohort. M2 macrophages are anti-inflammatory and promote tissue repair and neuroprotection, in contrast to the proinflammatory action of M1 macrophages [81, 82]. Previously, we observed that circulating monocytes from the Western cohort exhibited upregulation of genes associated with M1 polarization, and a non-significant trend for downregulation of M2 [40]. Results from the present work are consistent with these findings, suggesting that the Western cohort are polarized more towards a proinflammatory phenotype. This is further supported by the neuroanatomical data, as *MRC2* transcript levels, markers of anti-inflammatory M2 macrophages, were negatively correlated with total brain volumes, gray matter volumes, and temporoparietal volumes and thicknesses, and positively associated with white matter and CSF volumes.

In addition, we observed alterations in circulating monocyte expression profiles which were correlated with transcript levels in the temporal cortex and consistent with anti-inflammatory effects of the Mediterranean Diet relative to the Western Diet, and with the hypothesis that peripheral inflammation may promote neuroinflammation. Thus, potential interventions to increase anti-inflammatory activity may have a role in prevention or treatment of systemic as well as neuroinflammation.

*SLC3A2* was also lower in the Western cohort and showed negative relationships with time spent alone and percent change in cortical thickness, and positive associations with white matter volume. *SLC3A2*, which encodes SLC3A2 (the heavy subunit of cluster of differentiation 98, CD98), is an amino acid transporter and anti-apoptotic factor [83]. Although SLC3A2 and CD98 exhibit a complex relationship with inflammation (e.g., upregulated in intestinal epithelium in Crohn’s disease [84]), there is evidence that the action of this gene is downregulated in the CNS during proinflammatory states. In a study of microglia-derived extracellular vesicles, CD98 was present in control vesicles (i.e., those derived from un-activated microglia) but not in experimentally activated microglia, suggesting downregulation in the proinflammatory state [85].

Three other transcripts which were significantly reduced in the temporal cortex of the Western diet consumers have no previous published direct associations with neuropathology. Butyrophilin subfamily 2 member A1 (*BTN2A1*) encodes an integral plasma membrane protein which plays a role in immunomodulation of γδ T-cells which have been shown to have important anti-neuroinflammatory and neuroprotective effects[86, 87]. Katanin regulatory subunit B1 (*KATNB1*) is a subcomponent of the katanin heterodimer which functions in the cleavage and disassembly of microtubules, and perhaps plays a role in microtubule homeostasis [88], axonal growth, and neuronal migration [89]. Transmembrane protein 268 (*TMEM268*) is a relatively understudied transmembrane protein overexpressed in brain which appears to be important for cell adhesion, proliferation, and viability. TMEM268 has been shown to interact with integrin subunit β4 (ITGB4) [90].

Our exploratory analyses of potential pathways yielded further evidence of biological networks associated with inflammation. Among the top canonical pathways identified by Ingenuity Pathway Analysis (IPA) were phosphatase and tensin homolog (PTEN) signaling as well as ciliary neurotrophic factor (CNTF) signaling, both of which exert distinct immunomodulatory (including inflammatory) effects [91–93] and which have been found to be associated with AD-related neuropathology [94, 95]. Furthermore, among the top master regulators identified by IPA were transcription factor Sp3 and chymase 1, both of which have been implicated in inflammatory regulation [96, 97] and are present in higher levels in the brains of AD patients [98, 99]. Finally, transforming growth factor beta 1 (TGFB1), which was reduced in the Western cohort, is a pleiotropic cytokine with multifarious effects on both innate and adaptive immune function [100] and has been variously described as pro- and anti-inflammatory [101].

While our study provides compelling experimental evidence of diet-induced alterations in transcriptional regulation in the brain potentially associated with neuroinflammation, it also suggests important future directions for mechanistic research concerning neuroanatomical effects of diet composition. The potential mechanisms by which the Western diet promotes— and the Mediterranean diet limits—neuroinflammation are numerous. Metabolic and microbial disturbances induced by Western diet consumption likely promote inflammation [102, 103], while the Mediterranean diet’s antioxidant-rich composition likely limits inflammation [104]. Relatedly, Western diets perturb hypothalamic-pituitary-adrenal (HPA) axis physiology [33] which itself promotes pro-inflammatory responses [105] that may instigate a cascade of AD pathologic processes [106]. Pinpointing the key mechanisms by which diet impacts the brain will prove critical for developing effective interventions for populations that do not adhere to a Mediterranean diet.

In addition, as indicated by our prior work [34], diet-induced inflammation may be further compounded by social inequalities. In developed nations, individuals of low socioeconomic status are more likely to consume a stereotypical Western diet and experience greater psychological stress [107, 108], which may further exacerbate neurocognitive decline. This project focused on female macaques, as there is a paucity of data from female animal models[43], sex is known to affect neurodegenerative disease etiologies [109], and females are at higher risk of AD [110]. Consequently, data from males are necessary for fully understanding diet-induced neuroanatomical effects and associated molecular mechanisms. In addition, our transcriptional data are restricted to a single area of the brain at risk for the early manifestations of AD pathology. Additional studies are planned to explore other brain areas of importance to cognitive and other functions. Finally, whether these brain changes specifically impact cognitive function remains to be determined.

In conclusion, our results provide transcriptomic, neuroanatomical, and behavioral support for the premise that the Mediterranean diet exerts anti-inflammatory effects and preserves brain homeostasis and resilience compared to a Western diet. Increased cortical thickness and decreased white matter in the Western cohort, combined with transcriptomic signatures of inflammation and blood-brain barrier disruption, and behavioral correlates of neuroinflammation, suggest that the monkeys fed a Western diet developed neuroinflammation. The neuroanatomical variation we identified has important clinical relevance: increases in gray matter volume or cortical thickness associated with neuroinflammation could explain the progression of AD in the preclinical stage [36, 111] better than cortical atrophy. Future work integrating data from a variety of physiological systems (e.g., the gut microbiome, cardiovascular system, immune function, HPA physiology, metabolic health, cognitive function) will help map the causal pathways by which diet precipitates neuroinflammation.

The impact of diet composition on neuroinflammation in NHPs suggests that changes in diet composition could be an effective intervention in humans to reduce the risk of neurodegenerative processes in aging. Population level diet modification in humans has been shown to be feasible as evidenced by 1) the National Cholesterol Education Program which led to decreases in dietary fat and cholesterol intake and circulating cholesterol concentrations [112], and 2) the FDA requirement to list trans fats on food labels and reduce trans fats in foods which reduced circulating levels of trans fats by more than 50% [113]. Our findings suggest that population-wide adoption of a Mediterranean-like diet pattern may provide a cost-effective intervention on brain aging with the potential for widespread efficacy.

## Consent Statement

not applicable

