## Supplemental Methods for "Mediterranean Diet Protects Against a Neuroinflammatory Cortical Transcriptome: Associations with Brain Volumetrics, Peripheral Inflammation, Social Isolation and Anxiety"

*Cell Type Deconvolution*

To determine whether diet-based differential gene expression was driven by variation in cell types sampled for bulk RNA-seq, we estimated cell type proportions for each sample using the function *brainCells* in R package "BRETIGEA" [1]. The *brainCells* function returns a "surrogate proportion" variable, which indicates whether each cell type is over- or under-represented within a given sample. Cell type distributions were estimated using the programmers' top 50 marker genes for each of six cell types: Astrocytes, endothelial cells, microglia, neurons, oligodendrocytes, and oligodendrocyte precursor cells. We used the *adjustBrainCells* function to adjust transcript levels for variation in cell type proportions. We subsequently tested for diet-based differences in cell type proportions and adjusted transcript levels using Wilcoxon signed-rank tests (*wilcox.test* function in R).

*Magnetic Resonance Imaging*

Magnetic resonance imaging (MRI) scanning was performed at the MRI facility at Wake Forest University School of Medicine. Scans were conducted on sedated monkeys (1) during the baseline phase and (2) after 31 months of the experimental phase using a 3T Siemens Skyra scanner (Siemens, Erlangen, Germany) and a circularly polarized, 32-channel head coil (Litzcage, Doty Scientific, SC) to obtain T1-weighted anatomic images. After aligning baseline scans to the UNC Primate Atlas [2], we used the Advanced Normalization Tools (ANTs) antsMultivariateTemplateConstruction.sh command to create a whole-head, study-specific template. We then used the ANTs nonlinear registration algorithm (antsRegistrationSyN.sh [3, 4]) to calculate longitudinal transformations within individuals. Probabilistic tissue maps were used to automatically segment the brain into the three main tissue types: gray matter, white matter, and cerebrospinal fluid. We then determined volumes for the following regions of interest (ROIs): total gray matter (tGM), cortical gray matter (cGM), white matter (WM), cerebrospinal fluid (CSF), and total brain volume (TBV). The cGM ROI represents gray matter in the prefrontal, frontal, parietal, insular, cingulate, temporal, and occipital lobes [2]. The total GM was calculated as the sum of the cortical and deep GM ROIs. The dGM ROI represents gray matter in the hippocampus, amygdala, caudate and putamen. We summed tGM and WM to generate TBV.

Using a previously described method [5], we generated volume- and thickness-based AD temporoparietal meta-ROIs using the AD clinical literature as a template [6-8]. Briefly, the cortical thicknesses were determined using the "*KellyKapowski*" tool included with ANTs 2.2.0 [9] with the native spaced T1w MR image and the cortical GM tissue segmentation. Each meta-ROI represented the following ROIs (as illustrated in Fig. 1): the angular gyrus, the inferior, middle, and superior temporal gyrus, entorhinal cortex, fusiform cortex, supramarginal gyrus, precuneus, and parahippocampus. These measures are distinct as cortical thicknesses exclusively reflect gray matter, whereas cortical ROI volumes reflect tissue from both gray and white matter. The thickness meta-ROI was calculated as the volume-weighted average of the thicknesses of the individual ROIs to take into account the overall size of individual regions. The volume meta-ROI was calculated as the sum of all the individual ROIs. We collected the mean cortical thickness values and mean volume of non-zero voxels in each meta-ROI using the FMRIB Software Library *fslstats* utility (University of Oxford, Oxford, UK)[10].

*Correlations between Gene Expression and Brain Volumes*

We assessed relationships between transcript levels from genes that were differentially expressed by diet and MRI measures. We first normalized transcript levels using the trimmed mean of M-values (TMM) method [11], which uses a normalization factor calculated from a weighted average that excludes genes with high and highly variable transcript levels. One sample is selected as the reference, and a normalization factor is generated for each of the remaining samples [11]. Following normalization, we used the *corr.test* function in R to analyze transcript levels against percent change in the following global brain volume measures: TBV, tGM, cGM, WM, and CSF. We also determined correlations with the four following temporoparietal meta-ROIs: Right and left meta-ROI cortical thickness, and right and left cortical volume. We limited meta-ROIs to temporal and parietal targets because they were spatially proximate to the region from which we measured gene expression.
